# Maternal control of early life history traits affects overwinter survival and seedling phenotypes in sunflower (*Helianthus annuus* L.)

**DOI:** 10.1101/2020.07.24.220517

**Authors:** Fernando Hernández, Roman B. Vercellino, Ignacio Fanna, Alejandro Presotto

**Author notes:** Correspondence authors: Fernando Hernández, Departamento de Agronomía, Universidad Nacional del Sur, San Andrés 800, 8000 Bahía Blanca, Argentina., Alejandro Presotto, Departamento de Agronomía, Universidad Nacional del Sur, San Andrés 800, 8000 Bahía Blanca, Argentina.

## Abstract

When cultivated and wild plants hybridize, hybrids often show intermediate phenotypic traits relative to their parents, which make them unfit in natural environments. However, maternal genetic effects may affect the outcome of hybridization by controlling the expression of the earliest life history traits. Here, using wild, cultivated, and reciprocal crop-wild sunflower (*Helianthus annuus* L.) hybrids, we evaluated the maternal effects on emergence timing and seedling establishment in the field, and on seedling traits under controlled conditions. In the field, we evaluated reciprocal crop-wild hybrids between two wild populations with contrasting dormancy (the high dormant BAR and the low dormant DIA) and one cultivar (CROP) with low dormancy. Under controlled conditions, we evaluated reciprocal crop-wild hybrids between two wild populations (BAR and RCU) and one CROP under three contrasting temperature treatments. In the field, BAR overwintered as dormant seeds whereas DIA and CROP showed high autumn emergence (∼50% of planted seeds), resulting in differential overwinter survival and seedling establishment in the spring. Reciprocal crop-wild hybrids resembled their female parents in emergence timing and success of seedling establishment. Under controlled conditions, we observed large maternal effects on most seedling traits across temperatures. Cotyledon size explained most of the variation in seedling traits, suggesting that the maternal effects on seed size have cascading effects on seedling traits. Maternal effects on early life history traits affect early plant survival and phenotypic variation of crop-wild hybrids, thus, they should be addressed in hybridization studies, especially those involving highly divergent parents like cultivated species and their wild ancestors.

## INTRODUCTION

Hybridization, understood as the reproduction between individuals from divergent genetic lineages (historically isolated populations or related species), has played an important role in the evolution of wild and cultivated plants (Ellstrand & Rieseberg 2016). When hybridization occurs, the hybrid individuals harbor alleles from both the parents, which may increase their fitness through hybrid vigor (van Kleunen *et al*. 2015), transgressive segregation (Rieseberg *et al*. 2003), or adaptive introgression (Rieseberg *et al*. 2007). Hybridization may facilitate the colonization of novel habitats on evolutionary (Rieseberg *et al*. 2003) and ecological (Campbell *et al*. 2009; Mitchell *et al*. 2019) timescales. However, it can also disrupt favorable gene combinations and epistatic gene interactions when populations are locally adapted, leading to the expression of non-adaptive traits and then to outbreeding depression (Hall & Willis 2006). For hybridization to occur, the flowering period of sexually compatible plants must coincide in space and time, and when that happens the probability that hybridization succeeds will depend on the relative fitness of the hybrids relative to the non-hybrid progeny (Mercer *et al*. 2007).

Despite that a lot of work has been dedicated to studying the ecological and evolutionary consequences of hybridization in plants, much less attention has been paid to how maternal parents influence the phenotypes of their offspring (i.e. maternal effects) immediately after hybridization (Kimball *et al*. 2008; Piskurewicz *et al*. 2016; Singh *et al*. 2017). The maternal effect, defined by Roach and Wulff (1987) as “the contribution of the maternal parent to the phenotype of its offspring beyond the equal chromosomal contribution expected from each parent”, can be arbitrarily split into two groups: i) maternal environment effects, where phenotypic differences in the offspring are due to differential environmental conditions experienced by the maternal parents (Chiang *et al*. 2011; Fernández Farnocchia *et al*. 2019); and ii) maternal genetic effects, where phenotypic differences in the offspring are due to the maternal parent’s phenotype or genotype (Roach & Wulff 1987; Pace *et al*. 2015; Singh *et al*. 2017). Here, we focus on maternal genetic effects (hereafter simply referred to as maternal effect) to study how the maternal phenotype controls the phenotypes of reciprocal crop-wild sunflower (*Helianthus annuus* L.) hybrids.

Maternal effects may be attributed to the maternal inheritance of plastids and seed coats, double doses of maternal alleles in the endosperm, and the effects of maternal provisioning (nutrients, hormones, proteins, transcripts) during seed development (Roach & Wulff 1987; Weiss *et al*. 2013). In addition, recent studies demonstrated that maternal effects can be mediated by precise mechanisms of gene imprinting, where the maternal allele of functional genes is preferentially expressed over the paternal allele, thus conferring the maternal phenotype to the offspring (Rodrigues & Zilberman 2015; Piskurewicz *et al*. 2016; Iwasaki *et al*. 2019). When the maternal parent is locally adapted, it is expected that maternal effects on the early life history traits facilitate hybridization by increasing the survival of the hybrid offspring (Kimball *et al*. 2008).

The crop-wild complex represents an excellent model system to study post-hybridization maternal effects. Due to domestication and modern breeding, cultivated and wild populations often present large differences in ecologically relevant traits, such as seed size and dormancy, seedling size, and flowering time (Snow *et al*. 1998; Campbell *et al*. 2009; Hernández *et al*. 2017). Moreover, some of these traits are key to survival without human intervention. For example, the timing of emergence is critical for survival and maximizing fitness, and seed dormancy is the main mechanism involved in its regulation (Donohue *et al*. 2010; Fernández Farnocchia *et al*. 2019). Wild populations have complex mechanisms to synchronize germination with the best environmental conditions, which maximizes survival at this stage, but also influences the timing of later phenological phases, such as flowering and seed dispersal (Donohue 2014). On the contrary, crops often lack these complex mechanisms, which may result in high maladaptive germination and a low probability of forming self-perpetuating populations (Dong *et al*. 2011; Weiss *et al*. 2013; Pace *et al*. 2015). On the other hand, crops may have beneficial traits, such as large seedlings (Campbell *et al*. 2009; Singh *et al*. 2017), high competitive ability (Mercer *et al*. 2007), and herbicide resistance (Merotto *et al*. 2016), which may increase hybrid fitness in the presence of intraspecific and/or interspecific competition or herbicide applications (Mercer *et al*. 2007).

Previously, we reported strong maternal effects on seed traits in reciprocal crop-wild sunflower hybrids (Hernández *et al*. 2017). We proposed that the large differences observed in the seed anatomy, morphology, and dormancy under controlled conditions between reciprocal crop-wild hybrids may have a direct impact on the timing of emergence in the field and probably indirect impacts on seedling size and performance. In this study, using wild, cultivated, and reciprocal crop-wild sunflower hybrids, we evaluated the maternal effects on emergence and successful seedling establishment in the field, and on seedling traits under a wide range of temperatures under controlled conditions. The specific aims were: i) to investigate whether the maternal effects on seed traits result in differential timing of emergence and successful establishment in reciprocal crop-wild hybrids; and ii) to explore the maternal effects on plant vegetative traits across three contrasting temperatures.

## MATERIALS AND METHODS

### Plant material

Reciprocal crosses between wild and cultivated *H. annuus* were used. Wild populations were collected in three contrasting environments from central Argentina (Cantamutto *et al*. 2010): Colonia Barón (BAR; S 36°10′, W 63° 53′), Diamante (DIA; S 32° 03′ S, W 60° 38′), and Rio Cuarto (RCU; S 33°09, W 64°20). Cultivated materials were represented by two commercial cultivars (F1 hybrids): Cacique CL (Criadero El Cencerro) and HS03 (Advanta Seeds) which were randomly selected. Reciprocal crosses were made between the wild populations and cultivars (referred to as CROP). The crosses (hereafter referred to as Biotypes) were characterized according to their female and male parents.

We performed two experiments aiming to evaluate maternal effects in the field (Experiment I) and in the growth chamber (Experiment II). To minimize the environmental effects, all the achenes (hereafter seeds) used for the experiments were produced in a common garden at the Agronomy Department, Universidad Nacional del Sur, Bahia Blanca, Argentina (S 38°41’38’’, W 62°14’53’’). For Experiments I and II, seeds were produced during the 2016/2017 and 2017/2018 growing seasons, respectively. The seeds were produced under controlled hand-pollination of heads from 10-15 plants covered with paper bags at the pre-flowering stage, following Presotto *et al*. (2014) and Hernández *et al*. (2017). At the flowering stage, heads from wild plants were pollinated with pollen from sibling plants (wild biotypes: BAR, DIA, and RCU) or pollen from cultivated plants (wild-crop hybrids: BAR x CROP, DIA x CROP, and RCU x CROP) 3-4 times per head. For producing crop-wild hybrid seeds, heads from CROP plants were emasculated daily in the morning and pollinated with the corresponding wild pollen (BAR, DIA, and RCU to produce CROP x BAR, CROP x DIA, and CROP x RCU, respectively). To produce seeds of the CROP biotype, heads were self-pollinated by covering the heads with paper bags at the R4 stage until harvest.

In experiment I, we used the wild populations BAR and DIA, two wild populations with contrasting seed dormancy (Presotto *et al*. 2014; Hernández *et al*. 2017), the HS03 cultivar and their reciprocal crop-wild hybrids. In experiment II, we used the wild populations BAR and RCU, the Cacique CL cultivar and their reciprocal crop-wild hybrids. Therefore, seven biotypes were evaluated per experiment (female parent at the left): BAR, BAR x CROP, CROP x BAR, and CROP in both experiments, DIA, DIA x CROP, and CROP x DIA in Experiment I, and RCU, RCU x CROP, and CROP x RCU in Experiment II.

### Experiment I: maternal effects on timing of emergence and overwinter survival in the field

For this experiment, seeds were collected in late March from a common garden and the biotypes were produced by hand-pollination, as explained above. Fresh seeds were sown on April 20 (during early autumn to simulate natural dispersal) and the experiment ended on October 24 (midspring). Seeds were sown in groups of 50 in 23 cm diameter plastic pots (10 L) in the field. The biotypes were arranged in a complete randomized design with six replicates. The number of seedlings and their phenological stages were monitored in each pot every 14-21 days throughout the experiment. We also measured, just before winter (July 13), the leaf width and length, and plant height in four representative plants per pot and we calculated leaf size as the product of leaf length and leaf width. Minimum and maximum temperatures were recorded daily at a nearby weather station (Davis Vantage Pro 2) at CCT-Bahía Blanca.

### Experiment II: maternal effects on seedlings across temperatures

In this experiment, we evaluated the maternal effects on early vegetative traits under three contrasting temperature treatments in a growing chamber. The three temperature treatments were (day/night): 15 / 10 °C, 22 / 18 °C, and 30 / 26 °C, with a 12 h photoperiod. Light was provided with eight fluorescent tubes (36-40 W). For each temperature, to break seed dormancy, seeds were placed in trays on wet filter paper, covered with transparent polyethylene bags and stored for 1 week at 5°C (ISTA, 2004). Germinated seeds were put into pots in a growth chamber. Each biotype was represented by five seedlings (one per pot), totaling 35 pots per temperature. Plants were kept in the growth chamber until they reached the 4-leaf stage. The design was a randomized complete block with five replicates, with the racks in the growth chamber as blocks. At the 2-leaf stage, we measured cotyledon width and length, leaf width and length, and the plant height in each plant. At the 4-leaf stage, we measured leaf width and length (in the first pair) and the plant height. At this stage, the plants were dissected into three parts: stem, leaves, and roots. The leaves were scanned to obtain leaf area and then dried at 60 °C for 7 days, while the roots and stems were dried immediately for 7 days, and then weighed to obtain the leaf biomass, root biomass, and aerial biomass (stem + leaves) for each plant. The leaf area of two pairs of leaves per plant was measured using the software ImageJ (Schneider *et al*. 2012), then the leaf area of the four leaves was added to estimate leaf area per plant.

### Statistical analysis

In Experiment I, to characterize the extent of maladaptive autumn emergence, seedling overwinter survival, and the number of successfully established seedlings, we compared biotypes at three different times, respectively: end of autumn (June 12; T1), end of winter (September 4; T2) and mid-spring (October 24; T3). For each time, we compared the proportion of seedlings alive relative to the number of planted seeds (50 per pot), hereafter seedlings pot^-1^. The analyses of variance of seedlings pot^-1^ were performed using generalized linear models with PROC GLM (SAS University edition; SAS Institute Inc., Cary, NC). Biotype effect was considered as fixed, when the Biotype effect was significant, pairwise comparisons between Biotypes were made using Tukey-Kramer adjustment for multiple comparisons.

To analyze the phenotypic variation of emerged seedlings, we performed a multivariate analysis of variance (MANOVA) using the four variables (Leaf length, Leaf width, Plant height and Leaf size) with PROC GLM (SAS University edition; SAS Institute Inc., Cary, NC) using the Wilks’ lambda test criterion. All variables were ln-transformed to improve homoscedasticity. Biotype and individuals nested within pot were considered as fixed. Once the overall phenotypic variation showed a significant Biotype effect, each variable was analyzed using linear mixed models with PROC MIXED (SAS University edition; SAS Institute Inc., Cary, NC). The biotype effect was considered as fixed and individuals nested within pot as random. For variables with a significant Biotype effect, the biotypes were compared using the Tukey-Kramer adjustment for multiple comparisons.

In Experiment II, we measured the plant height, cotyledon length and width, leaf length and width at the 2-leaf stage and we estimated the cotyledon and leaf size as the product of length and width. At the 4-leaf stage, we measured the leaf length and width, plant height, leaf area, and biomass of leaves, stems, and roots. Then, from these variables we estimated the leaf size, specific leaf area (leaf area per plant / leaf biomass per plant), leaf biomass, aerial biomass, total biomass, and three allocation variables: leaf / aerial biomass, leaf / total biomass, and aerial / total biomass. In total, 21 phenotypic variables were recorded for each plant. To avoid redundancy, we performed a multiple Pearson’s correlation analysis with the *rcorr* function using the R package *Hmisc* (Harrell 2020). Then, for all the pairwise comparisons with r ≥ |0.9|, we removed one variable of each pair. After this filtering, a set of 10 variables was retained: Cotyledon size (cm^2^), Leaf size (cm^2^) at the 2-leaf stage (Leaf size_1), Leaf size and Plant height (cm) at the 4-leaf stage (Leaf size_2 and Plant height_2, respectively), Root biomass (mg), Aerial biomass (mg), Specific leaf area (cm^2^ mg^-1^), and three allocation variables named as Leaf / Total biomass, Leaf / Aerial biomass, and Aerial / Total biomass (Table S1).

First, all variables were ln-transformed to improve homoscedasticity. MANOVA was performed using the 10 variables with PROC GLM (SAS University edition; SAS Institute Inc., Cary, NC) using the Wilks’ lambda test criterion. Biotype, Temperature, Biotype*Temperature, and Block nested within temperatures were considered as fixed. If the overall phenotypic variation showed significant effects of either Biotype or Temperature, or a Biotype*Temperature interaction, the 10 variables were analyzed using linear mixed models with PROC MIXED (SAS University edition; SAS Institute Inc., Cary, NC). Biotype, Temperature, and Biotype*Temperature interaction were considered as fixed effects, while Block nested within Temperature was considered as random. Pairwise comparisons between Temperatures and between Biotypes were made using the Tukey-Kramer adjustment for multiple comparisons.

## RESULTS

### Experiment I

During the experiment, the monthly temperatures and annual precipitation were within the historical weather averages, but 2017 was wetter during autumn and dryer during summer (Table S2). Nine days after sowing, there were 10 successive warm days (from April 29 to May 8) with TMEAN > 15°C, T_MIN_ > 8°C, and T_MAX_ > 23°C (Fig. 1A), only interrupted by a day with TMEAN = 14.6°C, T_MIN_ = 5.8°C, and T_MAX_ = 21°C (May 4; Fig. 1A). Therefore, the temperature and water requirements for seed germination were met during this 10-day period.

**Figure 1.**
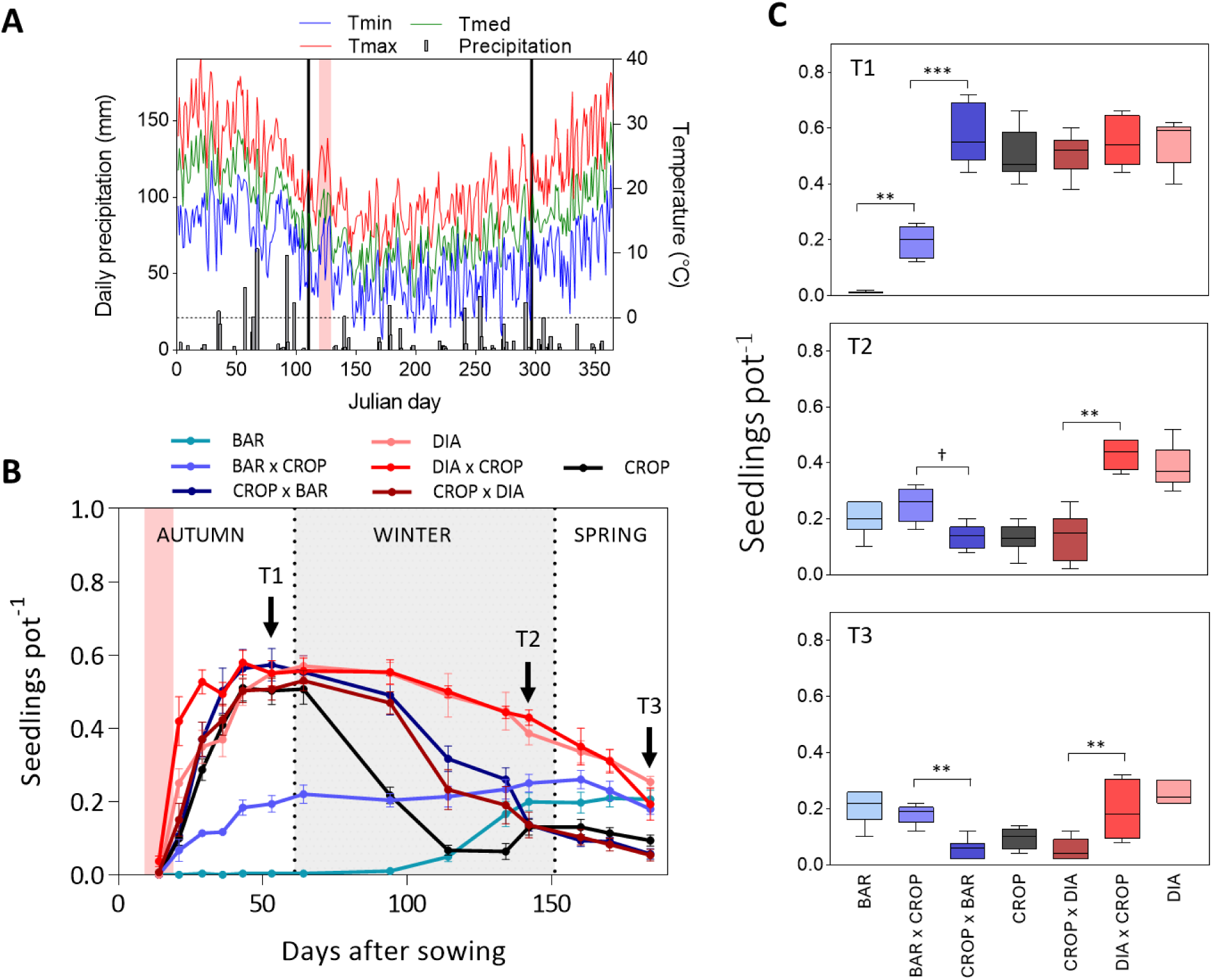
The timing of emergence in wild, cultivated, and crop-wild sunflower. A: Temperature variation and daily precipitation during 2017, vertical black lines indicate the beginning and the end of the experiment (from April 20 to October 24), and the red belt between 119 and 129 days (April 29 to May 8) indicates a 10-day period with warm days. B: Proportion of seedlings in each count relative to planted seeds (n = 50 per pot), red belt indicates the 10-day period with warm days. C: Biotype differences in the proportion of seedlings pot-1 at each time. Box edges represent the 0.25 and 0.75 quartiles, solid line represents the median value and error bars 0.1 and 0.9 percentiles. Outliers are shown. To facilitate interpretation, significant differences are only shown for the comparisons between biotypes with the same maternal parent and between reciprocal hybrids. *** P < 0.0001; **P < 0.01; *P < 0.05; † P < 0.1.

We observed an overall high emergence during autumn in DIA, CROP and their reciprocal crosses, with more than 50% of the planted seeds emerging during the first 45 days (Fig. 1B). This maladaptive autumn germination was negligible in BAR and low in BAR x CROP (Fig. 1B, 1C). On average, more than 50% of the seedlings died during winter (Fig. 1B) but differences in seedling survival between biotypes were evident (Fig. 1B). At the end of the experiment, the wild biotypes and crop-wild hybrids produced on wild plants had more established seedlings than CROP and crop-wild hybrids produced on CROP plants (Fig. 1) indicating a strong maternal effect on the number of established seedlings.

We found a significant Biotype effect at all three times (P < 0.0001). Among the biotypes, the proportions of seedlings relative to planted seeds (seedling pot^-1^) varied from 0 to 0.56 ± 0.04 seedlings pot^-1^ at T1, 0.14 ± 0.02 to 0.44 ± 0.02 seedlings pot^-1^ at T2, and 0.06 ± 0.02 to 0.26 ± 0.02 seedlings pot^-1^ at T3 (Fig. 1). At T1, BAR showed the lowest emergence, followed by BAR x CROP (Fig. 1B, 1C), whereas DIA, CROP, and their reciprocal hybrids showed more than 0.5 seedlings pot^-1^ (Fig. 1B, 1C). Significant differences were found between BAR and BAR x CROP (P = 0.0056), and between both BAR and BAR x CROP with the remaining biotypes (all P < 0.0001) while no significant differences were found for any comparison between DIA, CROP, and their reciprocal hybrids (all P > 0.7; Fig. 1C). When parents differed (i.e. BAR and CROP), the reciprocal hybrids showed more similarity to the female parent than to the male (Fig. 1) and significant differences were observed between them (BAR x CROP: 0.2 ± 0.02 seedlings pot^-1^, CROP x BAR: 0.58 ± 0.04 seedlings pot^-1^; P < 0.0001; Fig. 1C).

At T2, we evaluated the changes in the number of seedlings due to differential winter survival and new emergences during winter. DIA and DIA x CROP showed the highest values of seedlings pot^-1^ (0.38 ± 0.03 and 0.44 ± 0.02, respectively) due to a higher seedling survival during winter than in CROP, CROP x DIA, and CROP x BAR (Fig. 1B, 1C). No significant differences were observed between BAR and BAR x CROP (0.2 ± 0.02 and 0.26 ± 0.02, P = 0.8173), mostly due to the emergence during winter in BAR (Fig. 1B, 1C). No significant differences were found between biotypes with the same female parent (Fig. 1C). At T3, results were clear-cut, all the biotypes with wild maternal parents showed more seedlings pot^-1^ than the biotypes with crop maternal parents (mean wild maternal parents: 0.2 ± 0.02; mean crop maternal parents: 0.06 ± 0.01; Fig. 1B) and no significant differences were observed in comparisons within these two groups (Fig. 1B, 1C).

To explore the phenotypic variation in emerged plants, we measured the plant height, and leaf length and width on the same day (July 13) and we calculated the leaf size as the product of leaf length and leaf width. As BAR showed no emergence at that time, it was not included. In the MANOVA, we found significant phenotypic variation between the biotypes (F = 11.7; P < 0.0001) and no significant individual nested within pot effect (F = 1.2; P = 0.1669). For individual trait analyses, we found a significant Biotype effect in the four variables (Fig. 2). All traits showed maternal effects, and significant differences between reciprocal hybrids were observed in Leaf length, Plant height, and Leaf size for the hybrids between DIA and CROP, and in Leaf width, Plant height, and Leaf size for hybrids between BAR and CROP (Fig. 2).

**Figure 2.**
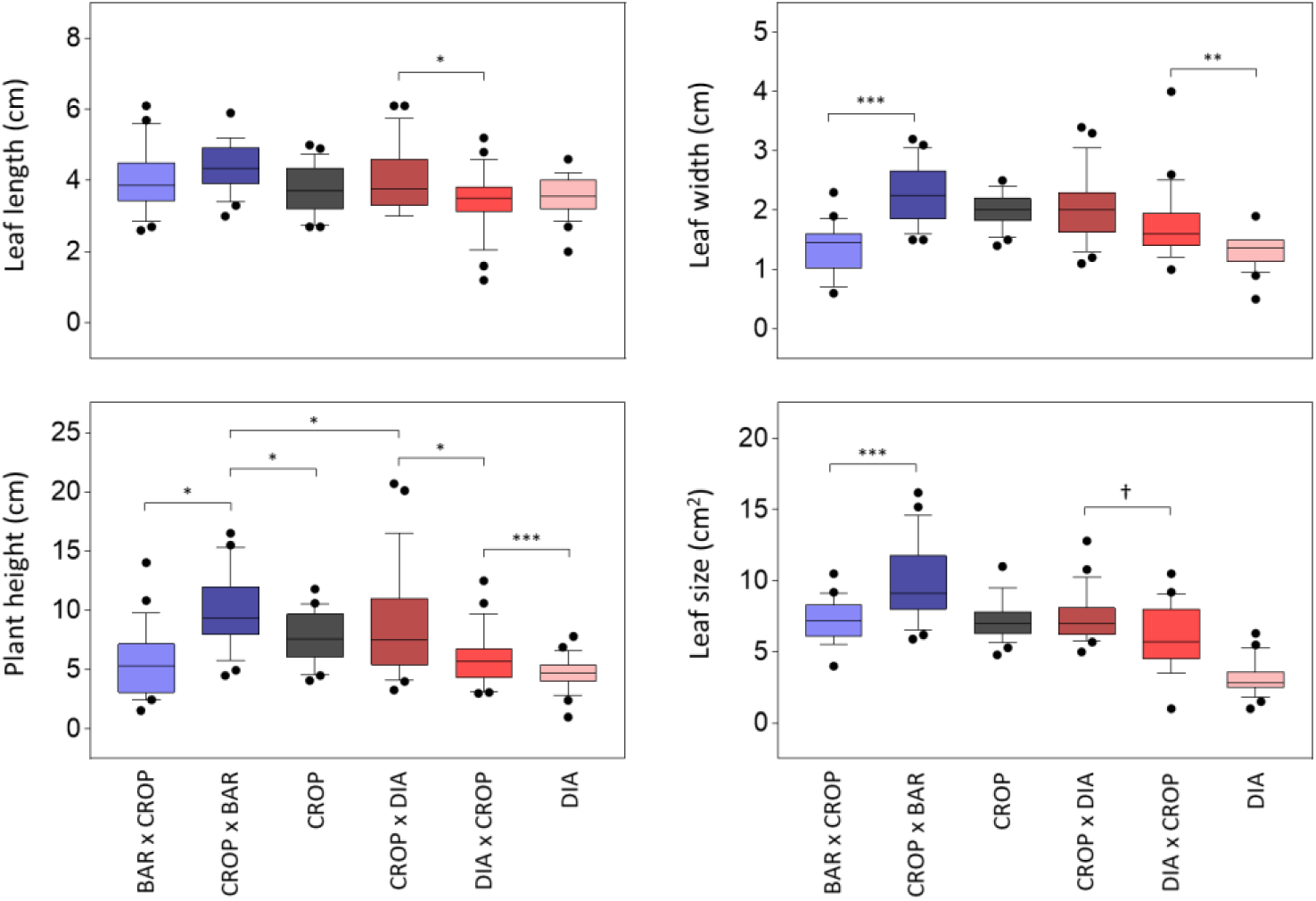
Phenotypic variation of seedling traits in the field. All variables on four representative plants per pot (24 plants per biotype) were recorded on the same day (July 13). All measured plants were at the 2-leaf developmental stage. Box edges represent the 0.25 and 0.75 quartiles, solid line represents the median value and error bars 0.1 and 0.9 percentiles. Outliers are shown. To facilitate interpretation, significant differences are only shown for comparisons between biotypes with the same maternal parent and between reciprocal hybrids. *** P < 0.0001; **P < 0.01; *P < 0.05; † P < 0.1.

### Experiment II

For the set of 10 variables, we found positive correlations between Cotyledon size and Leaf size (r = 0.73 and 0.72 for Leaf size_1 and Leaf size_2) and final biomass, both Aerial (r = 0.85) and Root biomass (r = 0.62), whereas Plant height showed a negative correlation with Leaf / Aerial biomass (r = -0.49) and a positive correlation with Aerial / Total biomass; r = 0.47). Specific leaf area showed negative correlations with most traits (r = 0.16 to -0.64) and, in general, the allocation traits showed positive correlations with Leaf size (Table S1).

In the MANOVA, we found significant effects of Temperature (F = 14.2; P < 0.0001), and Biotype (F = 2.8; P < 0.0001), but no significant Biotype*Temperature interactions (F = 1.0; P = 0.3682) or Block nested within temperatures effect (F = 0.9; P = 0.7278). Consistent with the MANOVA for individual trait analyses, we observed significant Temperature (in all the 10 traits), and Biotype (in eight out of 10 traits) effects but no significant Temperature*Biotype interaction for any trait, indicating that despite the large variation between temperatures observed for all traits (Fig. 3), the biotypes showed a similar response to temperatures. Under control conditions, the plants showed significantly higher Leaf size, lower Plant height, and they accumulated more Root biomass and Aerial biomass than the plants under low or high temperatures (Fig. 3). At low temperatures, plants showed larger cotyledons than plants under control and high temperatures and they showed higher Leaf size, Root biomass, and Aerial biomass than plants under high temperatures (Fig. 3). At high temperatures, the plants showed lower Cotyledon size and Leaf size (Fig. 3), and lower Root biomass and Aerial biomass than under control and low temperatures, but higher Plant height, and Specific leaf area than the plants under low and control temperatures (Fig. 3).

**Figure 3.**
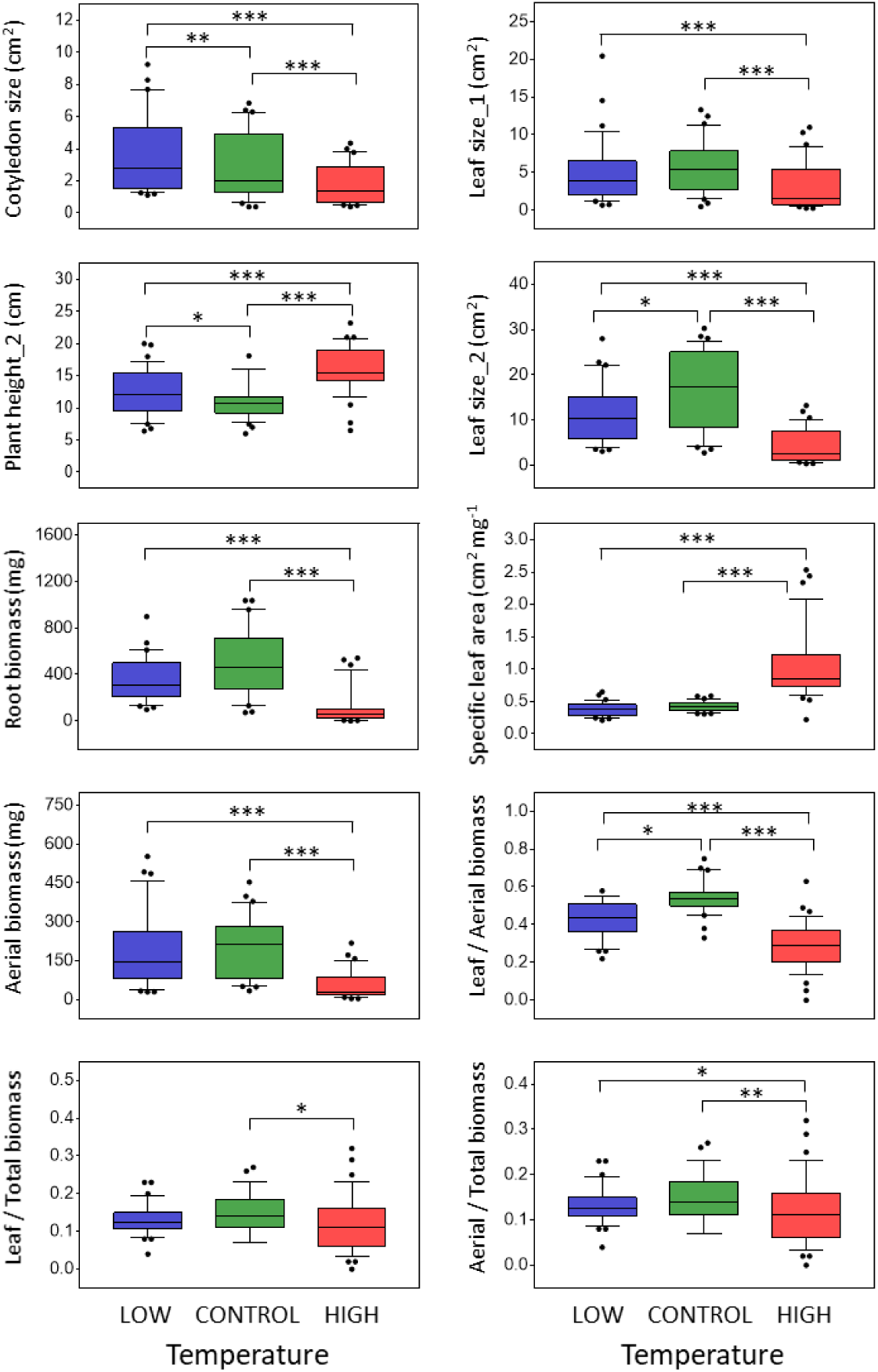
Phenotypic variation across three contrasting temperature treatments. Temperature treatments: LOW (15 / 10 °C), CONTROL (22 / 18 °C), and HIGH (30 / 26 °C) with neutral photoperiod (12 h light/12 h dark). Box edges represent the 0.25 and 0.75 quartiles, solid line represents the median value and error bars 0.1 and 0.9 percentiles. Outliers are shown. Significant differences between least square means of temperatures are indicated. *** P < 0.0001; **P < 0.01; *P < 0.05.

Significant differences were found between biotypes for eight out of 10 traits (all except Leaf / Aerial biomass and Leaf / Total biomass), and two of these traits (Specific leaf area and Aerial / Total biomass) showed no significant differences between either CROP and RCU or CROP and BAR. Thus, six traits were used to evaluate the maternal effects (Fig. 4). We observed strong maternal effects in all these six traits (Fig 4). Significant differences between CROP and the two wild populations (BAR and RCU) were observed in all traits (Fig. 4). CROP showed higher Cotyledon size, Leaf size, and Plant height (Fig. 4) than the wild populations, which resulted in higher Aerial biomass and Root biomass (Fig. 4). In addition, we observed no significant differences between BAR and RCU for any trait.

**Figure 4.**
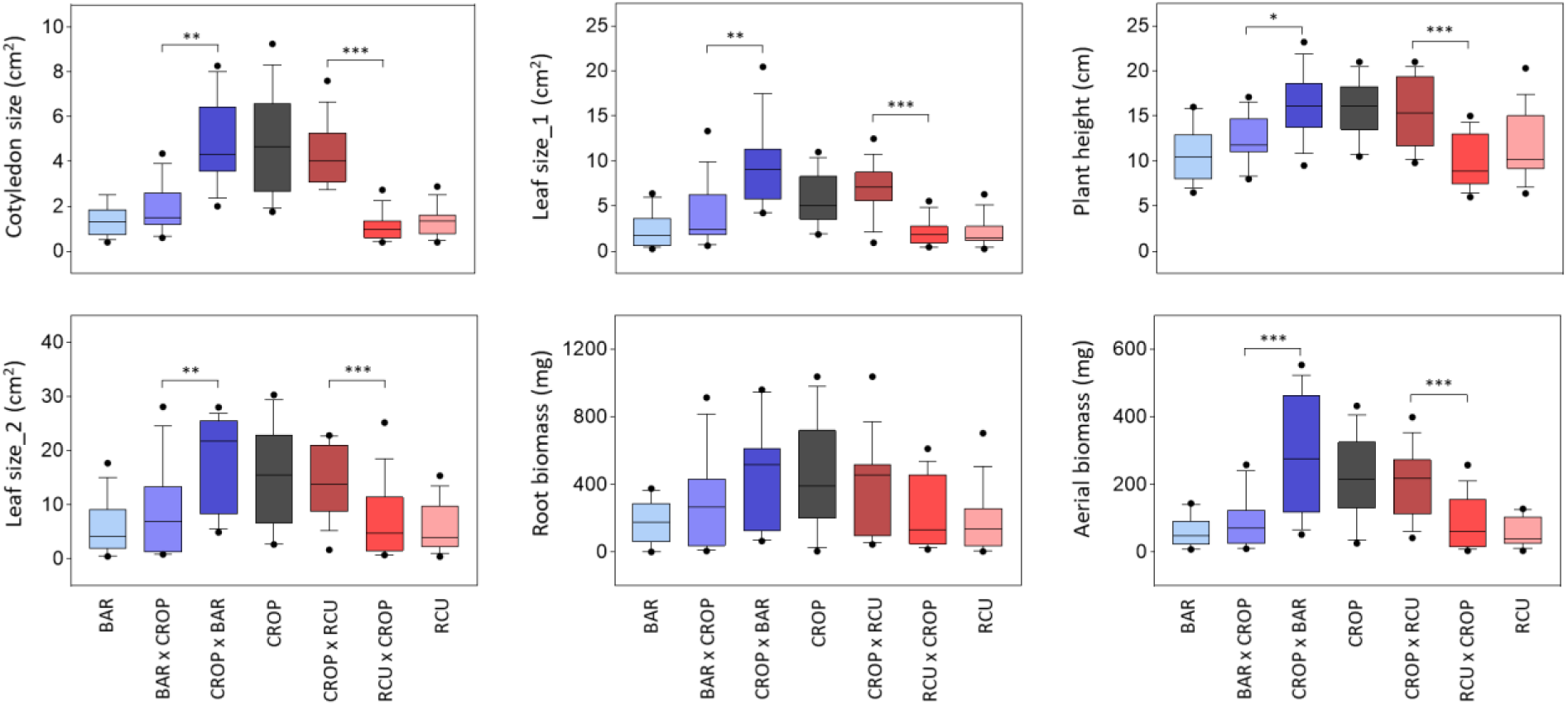
Phenotypic variation of six seedling traits across three temperature treatments. Box edges represent the 0.25 and 0.75 quartiles, solid line represents the median value and error bars 0.1 and 0.9 percentiles. Outliers are shown. To facilitate interpretation, significant differences are only shown for comparisons between biotypes with the same maternal parent and between reciprocal hybrids. *** P < 0.0001; **P < 0.01; *P < 0.05.

When the biotypes with the same maternal parent were compared, we observed no significant differences for any trait, i.e. crop-wild hybrids were statistically undistinguishable from their maternal wild or cultivated parents (Fig. 4). However, pairwise comparisons between reciprocal hybrids showed significant differences in all traits, except for Root biomass (Fig. 4).

Finally, we performed an analysis of covariance (ANCOVA) to evaluate whether the maternal effects on seedling growth can be explained by Cotyledon size, a maternally inherited trait. In the ANCOVA, we found a significant Temperature effect in all traits (P < 0.05), a significant covariate effect (P < 0.05) in four out of five traits (all but Aerial biomass), but no significant Biotype effect or Biotype*Temperature interaction for any trait (P > 0.05), indicating that most of the variation observed between biotypes, but not between temperatures, are explained by the maternally inherited cotyledon size.

## DISCUSSION

The timing of germination is a precise mechanism of habitat selection, which determines the environment that plants experience throughout their life cycle, and it is mainly regulated by seed dormancy (Donohue 2014; Postma & Ågren 2016). Here, we demonstrated that the maternal genetic effects on seed dormancy largely determined the timing of emergence in reciprocal crop-wild hybrids in the field, exposing them to different environmental conditions and then influencing their successful establishment. We also observed that the maternal effects are expressed in the phenotypic traits of emerged seedlings and that this phenotypic variation probably contributes to overwinter seedling survival. In the growth chamber experiment, strong maternal effects were observed in traits related to morphology and growth in which wild and cultivated parents differed, e.g. leaf size, and biomass of roots and aerial parts. Our study makes two main contributions: firstly, due to differences in the timing of emergence between parents, reciprocal crop-wild hybrids experience different environmental conditions, and therefore different selection pressures immediately after hybridization. It largely confirms our predictions based on germination experiments of the ecological and evolutionary importance of the maternal effects on seed traits (Hernández *et al*. 2017). Secondly, the large maternal effects observed here on the complex quantitative traits related to morphology and growth of seedlings demonstrate that the maternal parents can influence the phenotypes of the offspring well beyond seed traits.

### Primary seed dormancy is key to preventing out-of-season germination

In the field experiment, we evaluated one crop cultivar with low dormancy and two wild populations with contrasting seed dormancy levels (Presotto *et al*. 2014; Hernández *et al*. 2017). In this study, after a 10-day period of optimal temperatures for germination, more than 50% of seeds of the low dormant biotypes (CROP and DIA) germinated and emerged, whereas no emergence was recorded in the high dormant BAR. This maladaptive autumn emergence condemned DIA and CROP plants to resist winter temperatures as seedlings or perish, while the high dormant BAR survived as dormant seeds. Such differences in autumn emergence between biotypes were predicted from germination experiments under controlled conditions (Presotto *et al*. 2014; Hernández *et al*. 2017), and they confirm that, in wild sunflower, primary seed dormancy is key to preventing out-of-season germination. Similar results have been reported for sunflower in the native area (Alexander *et al*. 2014; Kost *et al*. 2015), as wild populations showed high overwinter survival whereas most cultivated seeds did not overwinter, mostly due to high autumn germination and subsequent winter mortality.

In contrast, in a recent seed bank study, no differences in the timing of emergence were observed between the high dormant BAR and the low dormant CROP and DIA, all biotypes showed similar emergence timing during late winter (Presotto *et al*. 2020). Such differences may be attributable to the environmental conditions that seeds experienced after dispersal in the different experiments. Germination occurs when soil temperature and soil water potential match with the temperature and water potential required for germination, but such requirements depend on the level of dormancy of the seed, i.e. low dormant seeds can germinate in a wide range of temperatures whereas high dormant seeds need more specific conditions (Batlla & Benech-Arnold 2015). However, if the minimum conditions for seed germination are not met (e.g. temperatures below the base temperature for seed germination or low water potential in dry soils), both dormant and non-dormant seeds will remain in the soil until the environmental conditions become favorable for germination (Batlla & Benech-Arnold 2015; Finch-Savage & Footitt 2017). In our study, we demonstrated that the seeds found optimal thermal and water conditions to germinate during early autumn, resulting in high maladaptive germination of low dormant parents, but not in the high dormant ones. Thus, seed dormancy was key to preventing out-of-season germination.

### Maternal effects on seed dormancy regulate the timing of emergence in crop-wild hybrids

Besides genetic differences in emergence timing, we observed a strong maternal effect on this trait. When seeds were produced on CROP plants, more than 50% of seeds emerged in the first 45 days, regardless of the pollen source. Such maladaptive autumn emergence drastically decreased the fitness of crop-wild hybrids produced on CROP plants, and therefore the chances of crop-wild gene flow. However, when seeds were produced on BAR plants (the high dormant parent), crop hybridization increased seedling emergence relative to BAR. Similar increased germination with crop hybridization has been reported for many crop-wild complexes, including sunflower (Snow *et al*. 1998; Presotto *et al*. 2014; Pace *et al*. 2015), lettuce (Hooftman *et al*. 2007), and rice (Dong *et al*. 2011). In our study, the increased germination of BAR x CROP did not result in differential seedling establishment relative to BAR, however, under real situations with intraspecific and interspecific competition, early emergence may increase the competitive ability of hybrids (Snow *et al*. 1998; Kost *et al*. 2015; Gioria & Pyšek 2017). Similar results have been reported for crop-wild sunflower hybrids in the native area (Mercer *et al*. 2006; Alexander *et al*. 2014; Kost *et al*. 2015), as hybrids produced on wild plants showed high overwinter survival (due to maternal effects on seed dormancy), and rapid germination during early spring (due to nuclear crop alleles).

From our data, it seems clear that seed dormancy of maternal genotypes influences the overwintering ability of crop-wild hybrids. In wild sunflower, clear geographic patterns of seed dormancy have been reported for native and invasive areas (Snow *et al*. 1998; Hernández, Poverene, *et al*. 2019). This information is useful for better predicting the risk of crop to wild gene flow at a regional scale. However, several factors may affect our prediction ability. Firstly, cultivated lines may present genetic variation for primary dormancy (Mercer *et al*. 2006), affecting seed dormancy in crop-wild hybrids. In addition, in locations where locally adapted populations have low dormant phenotypes, it is probable that mechanisms other than seed dormancy are important for overwinter survival (Postma & Ågren 2016); e.g. in Diamante, Entre Rios, where DIA was collected, the occurrence of freezing temperatures is not common, so the low dormancy levels (Presotto *et al*. 2014; Hernández *et al*. 2017) along with the extended flowering period (Cantamutto *et al*. 2010; Hernández, Presotto, *et al*. 2019) probably result in unseasonal growth of this population. Therefore, seed bank experiments in local environments, where wild and cultivated populations grow in sympatry, are needed to better understand the prediction ability of seed traits on the risk of crop to wild gene flow.

### Maternal effects on seedling traits

Maternal genetic effects on early vegetative traits were recognized very early in the literature (Corey *et al*. 1976; Roach & Wulff 1987). However, their ecological and evolutionary implications have been surprisingly understudied. In the field, most seedlings that emerged in autumn died during winter, however, in the two parents with a high autumn emergence, DIA showed higher seedling survival than CROP, indicating differential overwintering ability as seedlings. Recently, higher freezing tolerance has been reported in wild than in cultivated sunflower under controlled and field conditions (Hernández *et al*. 2020), probably because of natural selection for early winter emergence in Argentina (Presotto *et al*. 2020). In reciprocal hybrids between CROP and DIA, we observed strong maternal effects on phenotypic traits and overwinter seedling survival. Hybrids produced on DIA plants survived 3.6-fold times higher than hybrids produced on CROP plants, suggesting that freezing tolerance is maternally inherited, and it may contribute to overwinter survival in crop-wild hybrids (Hernández *et al*. 2020). Similarly, differential seedling survival and fitness was observed in reciprocal hybrids between related species adapted to different altitudes (Campbell & Waser 2001; Kimball *et al*. 2008). Interestingly, Kimball *et al*. (2008) showed that F1 hybrids outperformed parental species in all the environments evaluated, but hybrids produced on locally adapted mothers outperformed their reciprocal, indicating that maternal effects for locally adapted traits and hybrid vigor may occur simultaneously.

In the growth chamber experiments, most measured traits related to morphology and growth were maternally inherited, but we found no support for maternal inheritance of allocation traits. Maternal effects on seedling traits have been reported using reciprocal crop-wild hybrids of *Phaseolus vulgaris* L. (Singh *et al*. 2017) and diallel crosses of *Arabidopsis thaliana* L. (Corey *et al*. 1976) and *Lupinus texensis* Hook (Helenurm & Schaal 1996), suggesting they are common in plants.

Concerning the mechanisms behind the maternal effects, Singh *et al*. (2017) proposed that most of the maternal effects on seedling growth are likely to be the result of the differential seed size and cotyledon reserves of the maternal parent. Similarly, Helenurm and Schaal (1996) observed that the maternal effects operated primarily through differences in the seed size of the parents (i.e. mothers with larger seeds produce larger seedlings), but many differences in reciprocal hybrids persisted after controlling for seed size, suggesting the presence of other mechanisms. In our study, the cotyledon size, which largely depends on the seed size, explained most of the variation in seedling traits between biotypes, indicating that, probably, the observed phenotypic variation in seedling traits is a by-product of the maternal effects on seed size. However, the maternal effects on early life history traits clearly have ecological and evolutionary consequences, as they affect the early plant survival of crop-wild hybrids. Thus, we propose that the maternal effects on early life history traits should be addressed in hybridization studies, especially those involving highly divergent parents like wild and domesticated ones.

## AKNOWLEDGMENTS

We thank the National Research Council of Argentina (CONICET) for a fellowship for FH, RBV and IF. We also thank to Iván Diez and Gonzalo Fernández-Reyes for their valuable field and laboratory assistance. This work was supported by the National Agency for Scientific and Technological Promotion (PICT 2012-2854) and by the Universidad Nacional del Sur (PGI 24/A204).

## SUPPLEMENTARY MATERIAL

Table S1. Phenotypic correlation coefficients between 21 traits across three temperatures at the individual plant level (n = 101).

Table S2. Monthly mean temperatures and precipitations in 2017 and its comparison with historical values (period 1981-2010).

