## SUPPLEMENTARY MATERIAL for "Maternal control of early life history traits affects overwinter survival and seedling phenotypes in sunflower (*Helianthus annuus* L.)"

Table S1. Phenotypic correlation coefficients between 21 traits across three temperatures at the individual plant level (n = 101). Higher correlations in green (positive) and red (negative) and lower correlations in white. In bold, the 10 variables retained for further analyses.

| | Traits | 1 | 2 | 3 | **4** | 5 | 6 | **7** | **8** | 9 | 10 | **11** | 12 | 13 | **14** | 15 | **16** | **17** | 18 | **19** | **20** | **21** |
|---|---|---|---|---|---|---|---|---|---|---|---|---|---|---|---|---|---|---|---|---|---|---|
| 1 | Plant height_1 | | ** | NS | * | NS | * | NS | *** | NS | NS | NS | NS | NS | NS | NS | NS | NS | NS | *** | NS | *** |
| 2 | Cotyledon length | 0.30 | | *** | *** | *** | *** | *** | *** | *** | *** | *** | *** | *** | *** | *** | *** | *** | *** | NS | * | NS |
| 3 | Cotyledon width | 0.17 | 0.90 | | *** | *** | *** | *** | ** | *** | *** | *** | *** | *** | *** | *** | *** | *** | *** | NS | NS | NS |
| **4** | **Cotyledon size** | 0.22 | 0.96 | 0.96 | | *** | *** | *** | ** | *** | *** | *** | *** | *** | *** | *** | *** | *** | *** | NS | NS | NS |
| 5 | Leaf length_1 | 0.05 | 0.67 | 0.69 | 0.67 | | *** | *** | NS | *** | *** | *** | *** | *** | *** | *** | *** | *** | *** | *** | ** | NS |
| 6 | Leaf width_1 | 0.23 | 0.78 | 0.74 | 0.75 | 0.89 | | *** | ** | *** | *** | *** | *** | *** | *** | *** | *** | *** | *** | ** | ** | NS |
| **7** | **Leaf size_1** | 0.14 | 0.73 | 0.70 | 0.73 | 0.93 | 0.96 | | * | *** | *** | *** | *** | *** | *** | *** | *** | *** | *** | ** | ** | NS |
| **8** | **Plant height_2** | 0.91 | 0.42 | 0.30 | 0.36 | 0.15 | 0.32 | 0.25 | | NS | NS | NS | NS | ** | NS | NS | NS | NS | NS | *** | NS | *** |
| 9 | Leaf length_2 | -0.18 | 0.66 | 0.74 | 0.70 | 0.83 | 0.73 | 0.73 | -0.10 | | *** | *** | *** | *** | *** | *** | *** | *** | *** | *** | ** | * |
| 10 | Leaf width_2 | 0.00 | 0.68 | 0.72 | 0.70 | 0.83 | 0.84 | 0.80 | 0.06 | 0.90 | | *** | *** | *** | *** | *** | *** | *** | *** | *** | ** | NS |
| **11** | **Leaf size_2** | -0.07 | 0.68 | 0.73 | 0.72 | 0.80 | 0.79 | 0.78 | 0.00 | 0.93 | 0.97 | | *** | *** | *** | *** | *** | *** | *** | *** | ** | NS |
| 12 | Leaf area | -0.01 | 0.66 | 0.65 | 0.66 | 0.83 | 0.84 | 0.84 | 0.05 | 0.84 | 0.91 | 0.92 | | *** | *** | *** | *** | *** | *** | *** | *** | NS |
| 13 | Stem biomass | 0.15 | 0.84 | 0.89 | 0.91 | 0.76 | 0.76 | 0.78 | 0.27 | 0.77 | 0.76 | 0.79 | 0.73 | | *** | *** | *** | *** | *** | NS | ** | NS |
| **14** | **Root biomass** | -0.10 | 0.55 | 0.62 | 0.62 | 0.61 | 0.59 | 0.60 | -0.04 | 0.71 | 0.72 | 0.75 | 0.74 | 0.66 | | *** | *** | *** | *** | *** | NS | *** |
| 15 | Leaf biomass | -0.09 | 0.65 | 0.69 | 0.70 | 0.80 | 0.77 | 0.82 | -0.01 | 0.85 | 0.87 | 0.92 | 0.93 | 0.83 | 0.77 | | *** | *** | *** | *** | *** | NS |
| **16** | **Specific leaf area** | 0.16 | -0.38 | -0.45 | -0.40 | -0.53 | -0.39 | -0.39 | 0.12 | -0.64 | -0.53 | -0.51 | -0.45 | -0.47 | -0.42 | -0.49 | | *** | *** | *** | *** | NS |
| **17** | **Aerial biomass** | 0.03 | 0.78 | 0.83 | 0.85 | 0.81 | 0.80 | 0.83 | 0.13 | 0.85 | 0.85 | 0.89 | 0.87 | 0.96 | 0.74 | 0.95 | -0.51 | | *** | ** | ** | NS |
| 18 | Total biomass | -0.08 | 0.64 | 0.71 | 0.71 | 0.70 | 0.68 | 0.70 | 0.00 | 0.79 | 0.79 | 0.83 | 0.82 | 0.78 | 0.98 | 0.86 | -0.47 | 0.85 | | *** | NS | ** |
| **19** | **Leaf / aerial biomass** | -0.49 | 0.09 | 0.13 | 0.09 | 0.47 | 0.31 | 0.33 | -0.49 | 0.63 | 0.54 | 0.54 | 0.55 | 0.18 | 0.46 | 0.53 | -0.66 | 0.36 | 0.46 | | *** | ** |
| **20** | **Leaf / total biomass** | 0.02 | 0.22 | 0.17 | 0.19 | 0.38 | 0.35 | 0.35 | 0.07 | 0.34 | 0.35 | 0.35 | 0.40 | 0.27 | -0.02 | 0.40 | -0.41 | 0.34 | 0.08 | 0.46 | | *** |
| **21** | **Aerial / total biomass** | 0.43 | 0.13 | 0.04 | 0.08 | -0.04 | 0.07 | 0.03 | 0.47 | -0.21 | -0.11 | -0.13 | -0.11 | 0.06 | -0.39 | -0.10 | 0.16 | -0.02 | -0.31 | -0.36 | 0.59 | |

*** P < 0.0001; **P < 0.01; *P < 0.05; NS P > 0.05.

Table S2. Monthly mean temperatures and precipitations in 2017 and its comparison with historical values (period 1981-2010). Numbers in red and blue indicate at least 10 mm below and above, respectively, the corresponding historical monthly precipitation value. Note that monthly temperatures were similar to historical values (< 2°C). Source of historical data: https://www.smn.gob.ar/estadisticas.

| Period | Montlhy mean temperature (ºC) | | | | | | | | | | | | Annual mean Temperature (°C) |
|---|---|---|---|---|---|---|---|---|---|---|---|---|---|
| | Jan | Feb | Mar | Apr | May | Jun | Jul | Aug | Sep | Oct | Nov | Dec | |
| Historical (1981-2010) | 23.5 | 22.5 | 20 | 15.5 | 11.5 | 8.5 | 7.9 | 10 | 12.5 | 15.5 | 18.5 | 21.5 | 15.6 |
| 2017 | 24.2 | 23.7 | 19.8 | 14.5 | 12.3 | 9.0 | 9.4 | 10.5 | 11.9 | 14.4 | 16.2 | 21.5 | 15.6 |
| | Monthly precipitation (mm) | | | | | | | | | | | | Annual precipitation (mm) |
| Historical (1981-2010) | 67.1 | 67.1 | 75.2 | 54.6 | 41.3 | 31.7 | 31.1 | 34.5 | 51.6 | 73.3 | 56.8 | 67.1 | 651.4 |
| 2017 | 13.2 | 86.2 | 110.2 | 99.8 | 39.6 | 52.9 | 20.0 | 43.9 | 57.3 | 70.3 | 47.3 | 33.2 | 673.9 |
